# Implications of cholesterol and sphingomyelin in STING phosphorylation by TBK1

**DOI:** 10.1101/2021.03.15.435550

**Authors:** Kanoko Takahashi, Takahiro Niki, Emari Ogawa, Kiku Fumika, Yu Nishioka, Masaaki Sawa, Hiroyuki Arai, Kojiro Mukai, Tomohiko Taguchi

**Author notes:** Correspondence and requests for materials should be addressed to K. Mukai and T. Taguchi.

## Abstract

Stimulator of interferon genes (STING) is essential for the type I interferon response induced by microbial DNA from virus or self-DNA from mitochondria/nuclei. In response to emergence of such DNAs in the cytosol, STING translocates from the endoplasmic reticulum (ER) to the Golgi, and activates TANK-binding kinase 1 (TBK1) at the trans-Golgi network (TGN). Activated TBK1 then phosphorylates STING at Ser365, generating an interferon regulatory factor 3 (IRF3)-docking site on STING. How this reaction proceeds specifically at the TGN remains poorly understood. Here we report a cell-free reaction in which endogenous STING is phosphorylated by TBK1. The reaction utilizes microsomal membrane fraction prepared from TBK1-knockout (KO) cells and recombinant TBK1. We observed agonist-, TBK1-, “ER-to-Golgi” traffic-, and palmitoylation-dependent phosphorylation of STING at Ser365, mirroring the nature of STING phosphorylation *in vivo.* Treating the microsomal membrane fraction with sphingomyelinase or methyl-β-cyclodextrin, an agent to extract cholesterol from membranes, suppressed the phosphorylation of STING by TBK1. Given the enrichment of sphingomyelin and cholesterol in the TGN, these results may provide the molecular basis underlying the specific phosphorylation reaction of STING at the TGN.

## Introduction

The detection of microbial pathogens with nucleic acid sensors is a typical principle of innate immunity^1,2^. DNA-sensing or RNA-sensing proteins localize at various subcellular compartments and, upon binding to foreign nucleic acids, trigger innate immune signalling for host defence^3^. The DNA-sensing nucleotidyl transferase cGAS^4^, its product cyclic GMP-AMP (cGAMP)^5^, and the cGAMP sensor STING^6^ (also known as MITA^7^, ERIS^8^, MPYS^9^, or TMEM173) comprise a critical cytosolic DNA-sensing pathway in mammalian cells. STING is postulated to act as a scaffold to activate the downstream protein kinase TBK1^6–8,10,11^. Activated TBK1 phosphorylates and activates IRF3, the essential transcription factor that drives type I interferon production^12^. During this process, TBK1 also phosphorylates STING at Ser365, generating the IRF3-docking site on STING^13^.

STING is an ER-localized transmembrane protein. Upon its binding to cGAMP, STING translocates immediately to the Golgi with COP-II vesicles^14,15^. We and others have shown that the COP-II-mediated translocation of STING from the ER is required to activate the downstream signalling pathway^14–17^. In two autoinflammatory diseases, STING-associated vasculopathy with onset in infancy (SAVI)^18,19^ and the COPA syndrome^20^, STING is constitutively activated without DNA stimulation and localize not to the ER but to the perinuclear compartments that include the Golgi^14,19,21–25^. The interaction between STING and the downstream kinase TBK1 requires the translocation of STING from the ER to the Golgi^14^. Phosphorylated TBK1, the active form of TBK1, is exclusively localized to a subdomain within the TGN^21^. Together, these results demonstrate that the Golgi is an organelle at which STING recruits and activates TBK1 for triggering the STING-dependent innate immune response^26^.

As to the molecular mechanism underlying STING activation at the Golgi/TGN, we provide the evidence that palmitoylation of STING at the Golgi is essential to activate the downstream signalling pathway^21,27,28^. Disturbing Golgi lipid order by treating cells with D-ceramide-C6, which is converted to short-chain sphingomyelin, suppresses the activation of STING pathway^21^. These results suggest that certain Golgi lipids are involved in the STING activation, besides STING palmitoylation. However, because of the technical difficulty to deplete these lipids in cells, their involvement in the activation of STING signalling pathway has not been addressed.

In this work, we report a cell-free reaction in which endogenous STING is phosphorylated at Ser365 by recombinant TBK1. With this reaction, we demonstrate that sphingomyelin and cholesterol in the Golgi membrane are critical for STING activation.

## Results

### A cell-free reaction in which endogenous STING is phosphorylated by recombinant TBK1

To directly assess the roles of Golgi lipids in STING activation, we sought to develop a cell-free assay in which endogenous STING is phosphorylated by TBK1. To eliminate any contribution of cytosolic/endogenous TBK1 to STING phosphorylation, we decided to use microsomal membrane fraction prepared from TBK1-deficient cells as a membrane source containing endogenous STING.

TBK1-KO mouse embryonic fibroblasts (MEFs) were generated using the CRISPR-Cas9 technology (Fig. 1a). We then validated the phosphorylation and membrane traffic of STING in these cells. TBK1-KO MEFs were stimulated with DMXAA, a membrane-permeable mouse-specific STING agonist. As expected, 60 min after DMXAA stimulation, we observed the phosphorylation of STING at Ser365 in parental wild-type (WT) MEFs (Fig. 1a). In contrast, the phosphorylation was mostly suppressed in TBK1-KO MEFs. These results suggested that TBK1 was a major kinase for STING at Ser365, consistently with the previous observation^13^. In both WT and TBK1-KO MEFs, 60 min after DMXAA stimulation, STING was translocated from the ER to the Golgi, as shown by the co-localization with GM130 (a cis-Golgi protein) and TGN46 (a TGN protein) (Fig. 1b).

**Figure 1.**
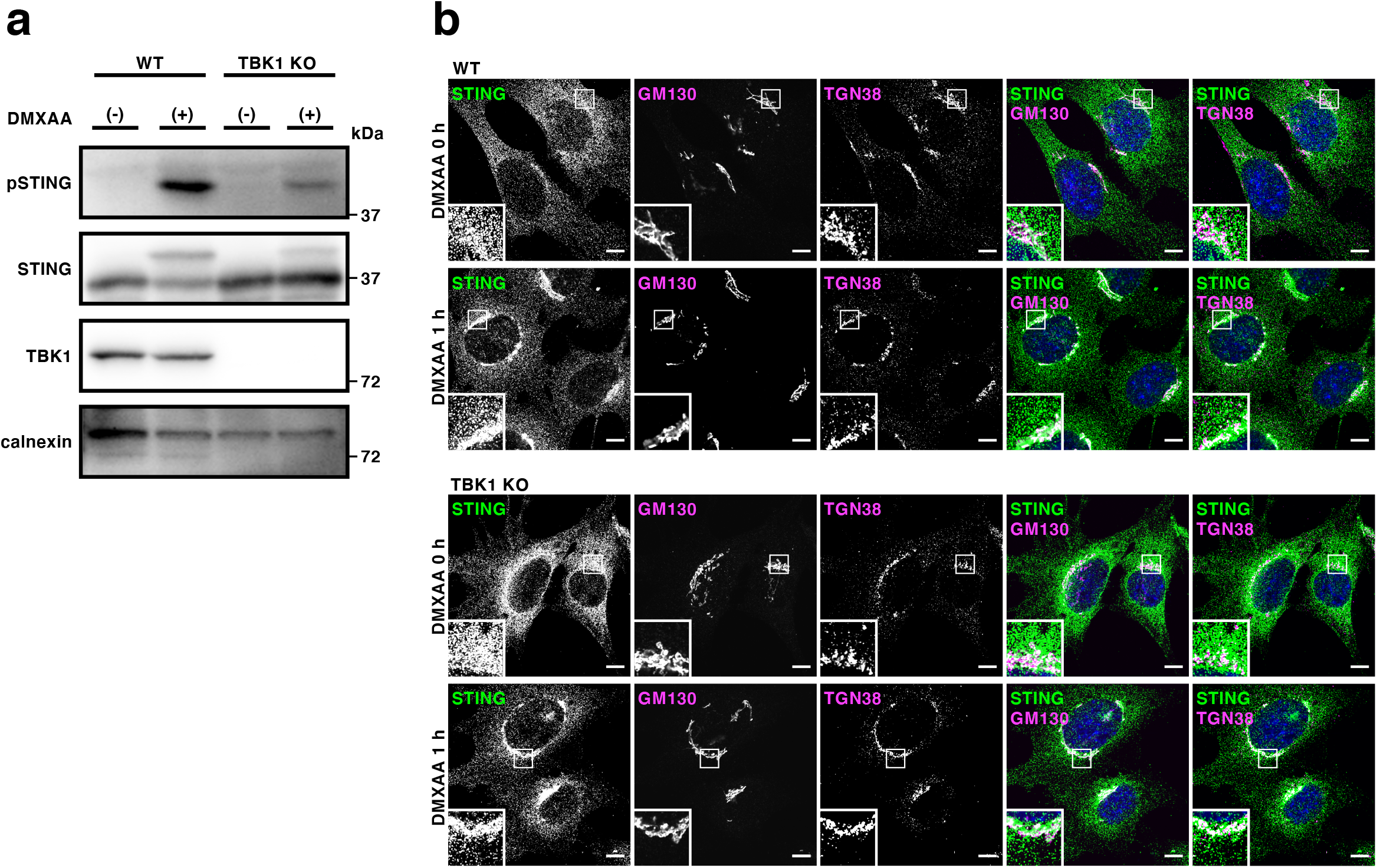
Generation of TBK1-knockout MEFs. (**a**) Western blots of cell lysates of WT or TBK1-knockout MEFs stimulated with DMXAA for 1 h. (**b**) Cells were stimulated with DMXAA for 1 h, fixed, permeabilized and stained for endogenous STING, GM130 (a cis-Golgi protein) or TGN38 (a trans-Golgi network protein). Nuclei were stained with DAPI (blue). Scale bars, 10 μm.

DMXAA-stimulated TBK1-KO MEFs were harvested in isotonic buffer and homogenized. The homogenates were then centrifuged at 3,000 x *g* for 5 min and the resulting post-nuclear supernatants were further centrifuged at 100,000 x *g* for 60 min.

The membrane fraction that floated just above 2 M sucrose cushion was collected and used as microsomal membrane fraction for the following *in vitro* reactions.

The microsomal membrane fraction was incubated with ATP and 100 ng of recombinant TBK1 at 37°C for the indicated time periods (Fig. 2a,b). In the conditions where microsomal membrane fraction prepared from DMXAA-stimulated MEFs was used, the amount of phosphorylated STING at Ser365 increased gradually up to 60 min of incubation (Fig. 2b). The amount of STING was decreased after 120 min of incubation, suggesting the proteolytic degradation of STING (Supplementary Fig. S1). When microsomal membrane fraction of unstimulated cells was used for the reaction, the phosphorylated STING disappeared (Fig. 2b). These results suggested that STING phosphorylation by TBK1 stringently requires the agonist (DMXAA) stimulation.

**Figure 2.**
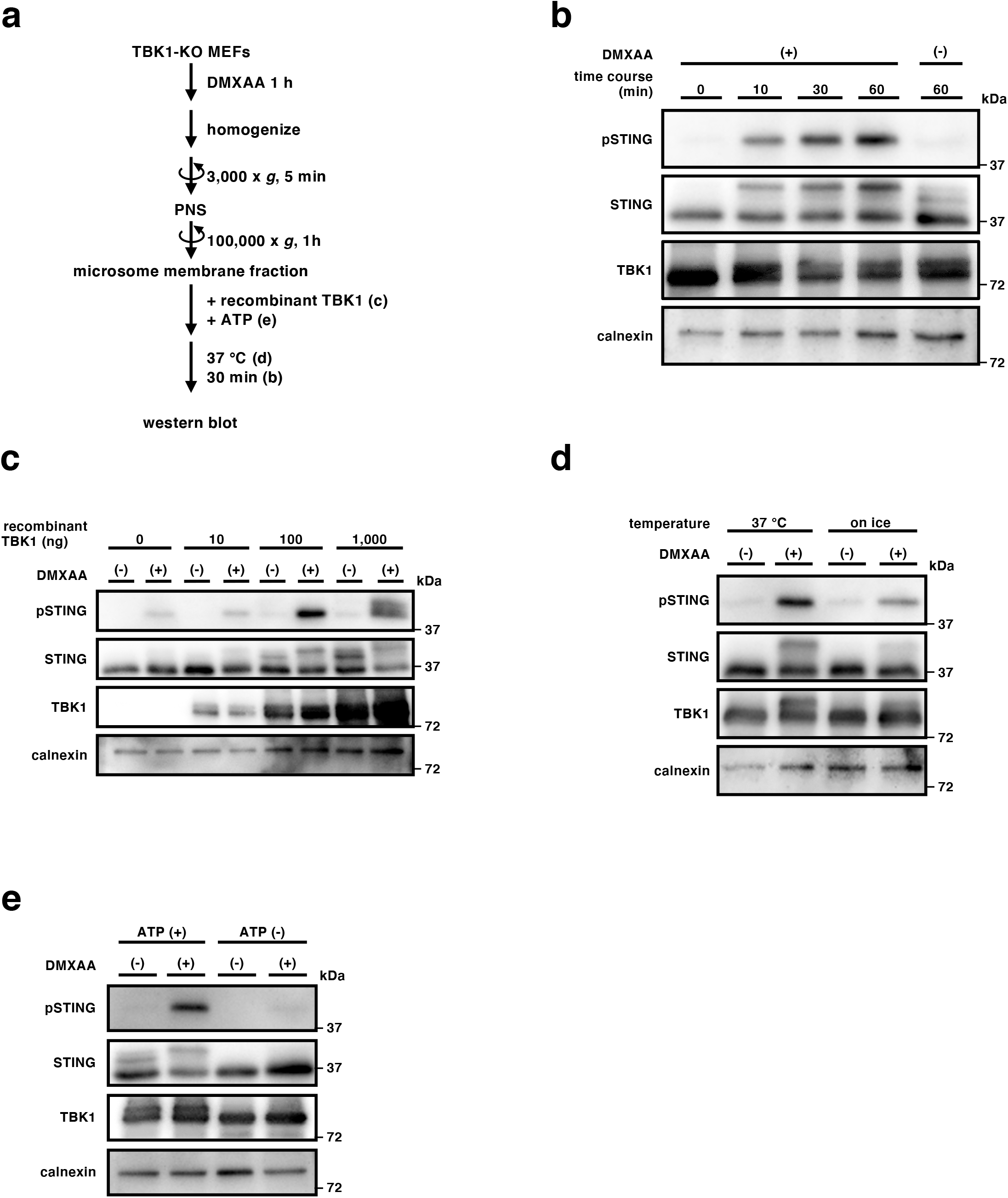
TBK1-dependent phosphorylation of STING *in vitro.* (**a**) A protocol for TBK1-dependent phosphorylation of STING *in vitro.* (**b**-**e**) TBK1-knockout MEFs were stimulated with DMXAA (25 μg/ml) for 0 or 1 h, homogenized in isotonic buffer, and centrifuged at 3,000 x *g* for 5 min. The resulting post-nuclear supernatants were then centrifuged at 100,000 x *g* for 1 h, and the pellets were resuspended in isotonic buffer. The resuspended membrane fractions were incubated with recombinant TBK1 (100 ng) and ATP (1 mM) at 37°C for 0, 10, 30, or 60 min (**b**). The membrane fractions were incubated with ATP and the indicated amount of recombinant TBK1 (0 ng, 10 ng, 100 ng or 1,000 ng) at 37°C for 30 min (**c**). The membrane fractions were incubated with ATP and recombinant TBK1 at 37°C or on ice for 30 min (**d**). The membrane fractions were incubated with recombinant TBK1 in the presence or absence of ATP at 37°C for 30 min (**e**). Phosphorylation of STING at Ser365 was then analyzed by western blot (**b-e**).

Titration of the amount of TBK1 in the reaction showed that 100 ng of recombinant TBK1 was sufficient to detect the phosphorylated STING (Fig. 2c). Thus, we routinely used 100 ng of recombinant TBK1 at the following experiments. The phosphorylation reaction of STING by TBK1 was temperature-sensitive (Fig. 2d) and required ATP (Fig. 2e). In sum, a cell-free assay to phosphorylate endogenous STING at Ser365 was developed with the use of recombinant TBK1.

### “ER-to-Golgi” traffic- and palmitoylation-dependent phosphorylation

The exocytic membrane traffic from the ER to the Golgi is required for the activation of the STING signalling pathway^14,21,22,29^. Thus, we examined if the phosphorylation of STING in the cell-free assay also required the translocation of STING to the Golgi.

Microsomal membrane fraction was prepared from cells treated with BFA (a fungal macrocyclic lactone that blocks “ER-to-Golgi” traffic^30^) and DMXAA. When the microsomal membrane fraction was subjected to the *in vitro* reaction, we found no phosphorylated STING (Fig. 3a). Addition of STING agonist *i.e.,* DMXAA or 2’3’-cGAMP, to the *in vitro* reaction with the microsomal membrane fraction prepared from unstimulated cells did not promote the phosphorylation of STING (Fig. 3b and Supplementary Fig. S2). These results suggested that (i) the binding of STING with its agonist alone was not sufficient to make STING the substrate for TBK1 and (ii) STING translocation from the ER to the Golgi, which subsequently occurred after the binding of STING with its agonist, was required for the phosphorylation of STING.

**Figure 3.**
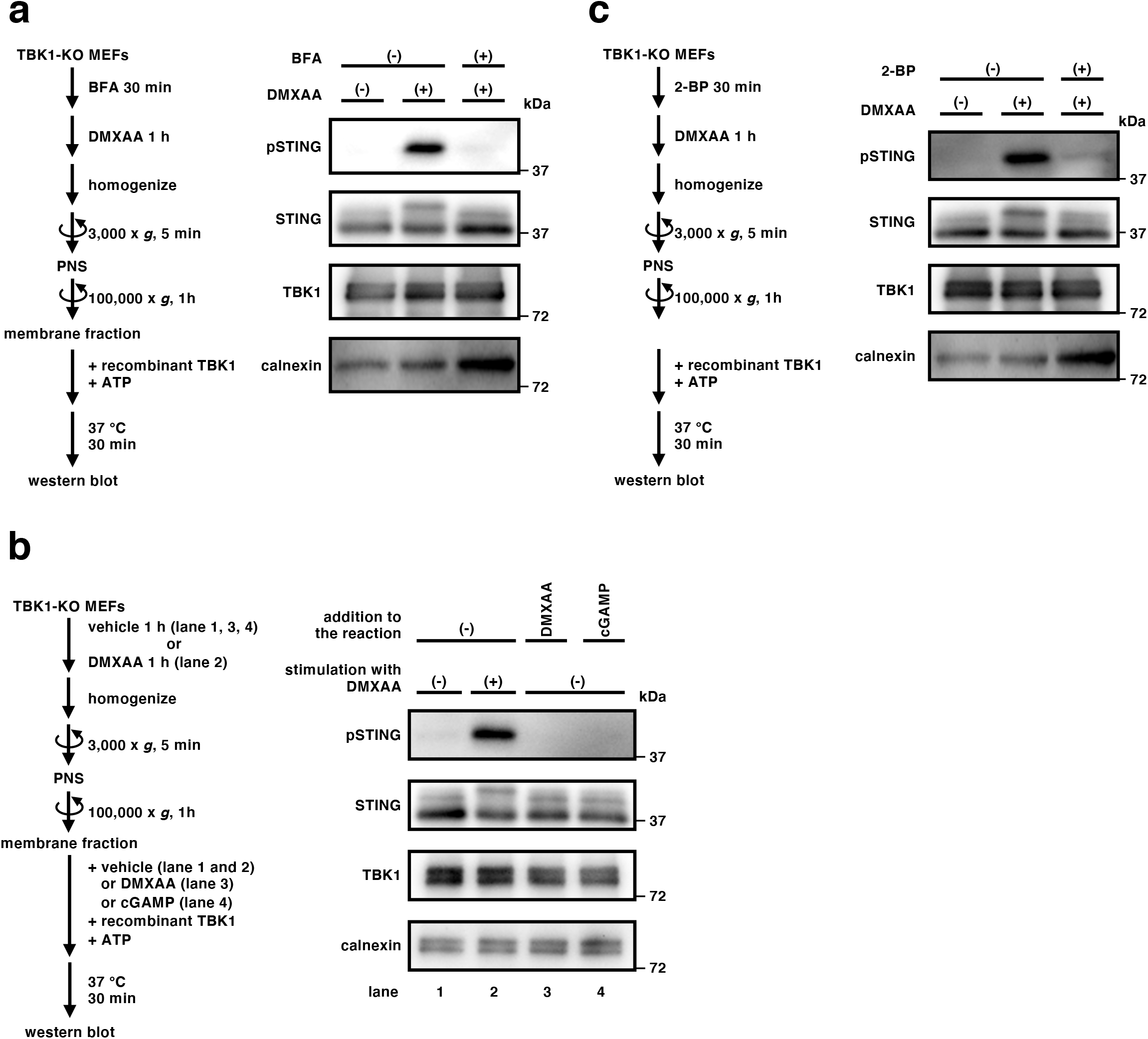
Phosphorylation of STING by TBK1 *in vitro* requires “ER-to-Golgi” traffic and palmitoylation of STING. (**a**) TBK1-knockout MEFs were treated with brefeldin A (3 μg/mL) for 30 min followed by the stimulation with DMXAA for 1 h. The post-nuclear supernatants of cells were centrifuged at 100,000 x *g.* The resulting membrane fraction was resuspended, incubated with recombinant TBK1 in the presence of ATP, and analyzed by western blot. (**b**) Post-nuclear supernatants of unstimulated TBK1-knockout MEFs were centrifuged at 100,000 x *g*. The resulting membrane fraction was resuspended and incubated with recombinant TBK1 and ATP in the presence of DMXAA (25 μg/ml) or 2’3’-cGAMP (250 ng/mL) at 37°C for 30 min. Phosphorylation of STING at Ser365 was analyzed by western blot. (**c**) TBK1-knockout MEFs were treated with 2-bromopalmitate (50 μM) for 30 min followed by the stimulation with DMXAA for 1 h. The membrane fraction of the cells was resuspended, incubated with recombinant TBK1 in the presence of ATP, and analyzed by western blot.

STING undergoes palmitoylation at the Golgi and the inhibition of palmitoylation with 2-bromopalmitate (2-BP) suppressed the STING-dependent downstream signalling^21^. Microsomal membrane fraction was prepared from cells treated with 2-BP and DMXAA. When the microsomal membrane fraction was subjected to the *in vitro* reaction, we found no phosphorylated STING (Fig. 3c), consistently with the *in vivo* observation^21^.

### A role of cholesterol and sphingomyelin in STING activation

We previously suggested the role of sphingomyelin in STING activation by the experiment with D-ceramide-C6^21^, an agent to disrupt lipid domain containing sphingomyelin^31^. We sought to address directly the role of lipids constituting the lipid domain in STING activation with the cell-free reaction. Microsomal membrane fraction prepared from DMXAA-stimulated cells was digested with recombinant bacterial sphingomyelinase (bSMase), and then subjected to the *in vitro* reaction. As shown in Fig. 4a, we found no phosphorylated STING with bSMase-digested membranes. Pretreatment of the microsomal membrane fraction with methyl-β-cyclodextrin, an agent to extract cholesterol from membranes, also suppressed the phosphorylation of STING by TBK1 in dose-dependent fashion (Fig. 4b). Given that STING mostly localized at the Golgi that includes the TGN 60 min after DMXAA stimulation^21^ (Fig. 1b), sphingomyelin and cholesterol in the TGN may participate in STING activation.

**Figure 4.**
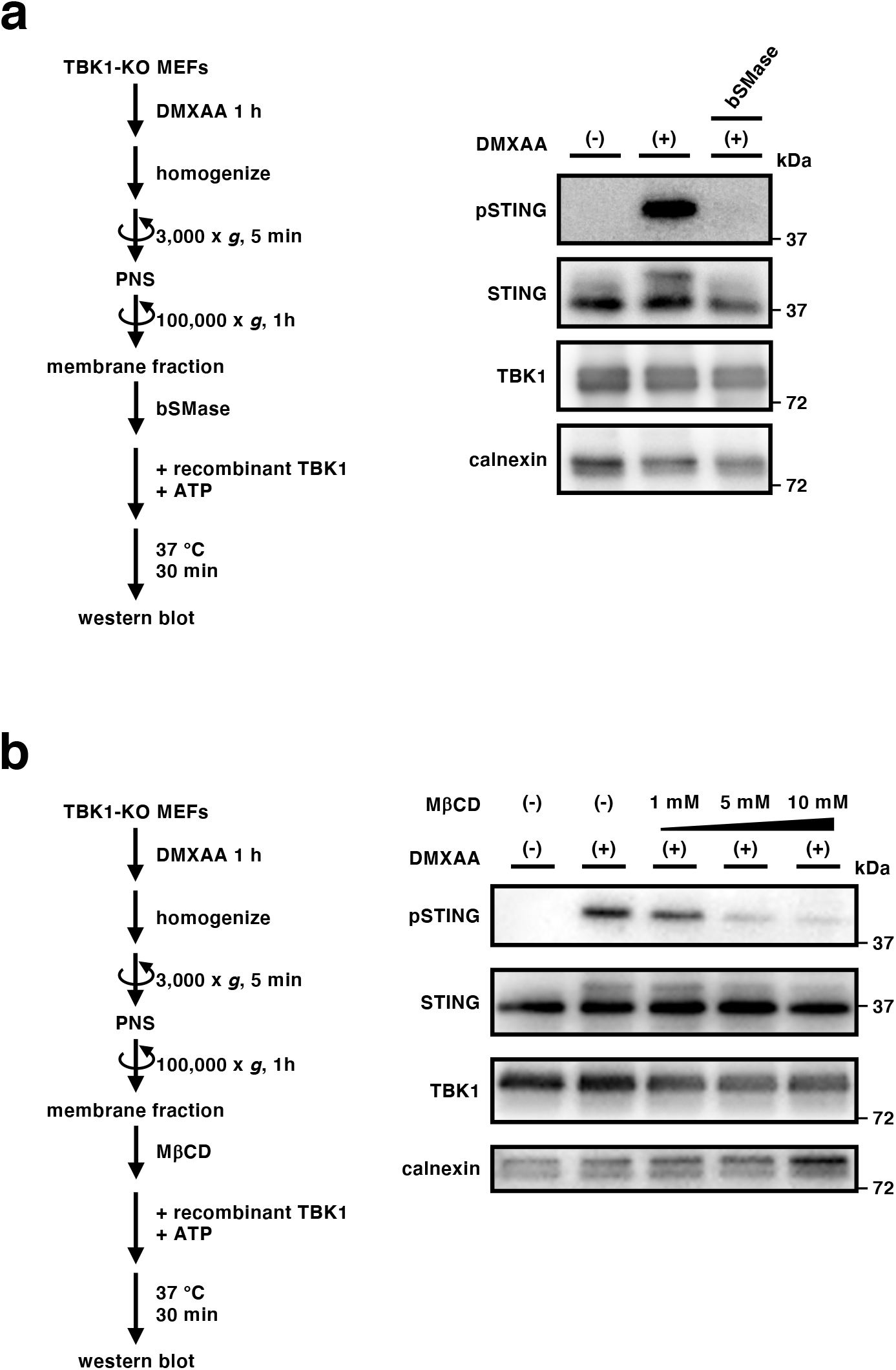
A role of cholesterol and sphingomyelin in STING phosphorylation *in vitro.* (**a-b**) The membrane fraction of TBK1-knockout MEFs that were stimulated with DMXAA for 1 h was collected by ultracentrifugation as in Figure 2. The resuspended membrane fraction was incubated with recombinant bacterial sphingomyelinase (0.125 U/mL) (**a**) or methyl-β-cyclodextrin (MβCD) (1 mM, 5 mM, or 10 mM) at 37°C for 30 min followed by incubation with recombinant TBK1 and ATP for 30 min. Phosphorylation of STING at Ser365 was then analyzed by western blot.

## Discussion

In the present study, we developed a cell-free assay in which endogenous STING is phosphorylated by recombinant TBK1. The assay showed the dependency of STING phosphorylation on the STING agonist, TBK1, “ER-to-Golgi” traffic, and palmitoylation, which mirrored the nature of STING phosphorylation *in vivo.* With the assay, we provided an *in vitro* evidence that sphingomyelin and cholesterol in the Golgi membranes were involved in the STING activation. The assay should provide a useful biochemical complement to cell biological studies presently used to understand the molecular mechanism underlying the STING activation.

Protein palmitoylation has been implicated in the clustering of a number of proteins into a specific lipid microdomain “raft” enriched in cholesterol and sphingomyelin^32^. Cholesterol is suggested to be enriched at the TGN, and cholesterol together with sphingomyelin generated by sphingomyelin synthase 1 are thought to form lipid rafts at the TGN^31^. Together with the present findings by the cell-free experiments, we hypothesize that palmitoylated STING becomes clustered at the TGN with the aid of the raft, which facilitates the recruitment of TBK1^10,11^ for phosphorylation of STING^13^.

Sphingomyelin at the TGN is demonstrated to be essential for transport carrier formation^31^. Cholesterol at the Golgi, the levels of which is regulated by oxysterol-binding protein, is essential for the Golgi localization of intra-Golgi v-SNAREs by ensuring proper COP-I vesicle transport^33^. Caveolin-1, a protein to form caveolae at the plasma membrane, accumulates in the Golgi when the cholesterol level in the Golgi is low^34^. The present study demonstrates another role of sphingomyelin and cholesterol at the Golgi/TGN in cellular signalling. The implication of these lipids in innate immune response may lead to new treatments for cytosolic DNA-triggered autoinflammatory diseases.

## Methods

### Antibodies

Antibodies used in this study were as follows: rabbit anti-TBK1 (ab40676, dilution 1:1000; Abcam); rabbit anti-phospho-STING (D8F4W, dilution 1:1000) (Cell signaling); rabbit anti-calnexin (10427-2-AP, dilution 1:1000), rabbit anti-STING antibody (19851-1-AP, dilution 1:1000 for western blot and immunofluorescence) (Proteintech); mouse anti-GM130 (610823, dilution 1:1000) (BD Biosciences); sheep anti-TGN38 (AHR499G, dilution 1:1000) (Bio-Rad); Goat Anti-Rabbit IgG(H+L) Mouse/Human ads-HRP (4050-05, dilution 1:10000) (Southern Biotech); Alexa 488-, 594-, or 647-conjugated secondary antibodies (A21203, A21206, A21448, dilution 1:2000) (Thermo Fisher Scientific)

### Reagents

The following reagents were purchased from the manufacturers as noted: DMXAA (14617, Cayman), brefeldin A (11861, Cayman), methyl-β-cyclodextrin (332615, Sigma), recombinant bacterial sphingomyelinase (S7651, Sigma); 2-bromopalmitate (320-76562, Wako), recombinant TBK1 (Carna Biosciences, Inc.). lentiCas9-Blast (Addgene plasmid # 52962) and lentiGuide-Puro (Addgene plasmid # 52963) were gifts from Feng Zhang. psPAX2 (Addgene plasmid # 12260) and pMD2.G (Addgene plasmid # 12259) were gifts from Didier Trono.

### Tag-free recombinant TBK1

N-Terminal GST-fusion full length human TBK1 protein (#05-115, Carna Biosciences) was treated with Turbo3C protease (Accelagen) at 4 °C overnight to cleave the GST-tag. The mixture was applied to a Glutathione Sepharose 4 Fast Flow column (#17-5132-03, Cytiva) and the flow-through fraction was collected.

### Cell culture

MEFs were obtained from embryos of wild-type C57BL/6JJcl mice at E13.5 and immortalized with SV40 Large T antigen. MEFs were cultured in DMEM supplemented with 10% fetal bovine serum and penicillin/streptomycin/glutamine in a 5% CO2 incubator.

### Generation of TBK1-KO cells by CRISPR-Cas9

MEFs that stably expressed Cas9 were established using lentivirus. HEK293T cells were co-transfected with lentiCas9-blast, psPAX2, and pMD2.G. The medium that contains the lentivirus was collected and filtered through 0.45 μm PVDF filter. WT MEFs were incubated with the medium for 6 h and then selected with blasticidin (5 μg/mL) for several days.

Single-guide RNAs (sgRNA) were designed to target mouse TBK1 genomic loci. The sgRNA (sense: 5’-caccgGGTGCACTATGCCGTTCTCT-3’, antisense: 5’-aaacAGAGAACGGCATAGTGCACCc-3’) was cloned into a lentiGuide-Puro vector. The lentiviral plasmids, psPAX2, and pMD2.G were then co-transfected into HEK293T cells and the lentivirus-containing medium was collected. Cas9-expressing MEFs were incubated with the medium for 6 h and selected with puromycin (2 μg/mL) for several days. Single colonies were then isolated and the expression of TBK1 in each clone was analyzed by western blot.

### Immunocytochemistry

Cells were fixed with 4% paraformaldehyde in PBS at room temperature for 15 min and permeabilized with digitonin (50 μg/mL) in 3% BSA-PBS at room temperature for 5 min. Cells were then incubated with primary antibodies, then with secondary antibodies conjugated with Alexa fluorophore.

### Confocal microscopy

Confocal microscopy was performed using a LSM880 with Airyscan (Zeiss) with a 63 × 1.4 Plan-Apochromat oil immersion lens. Images were analyzed with Zeiss ZEN 2.3 SP1 FP3 (black, 64 bit) (ver. 14.0.21.201) and Fiji (ver. 2.0.0/1.52n).

### *In vitro* assay of recombinant TBK1-dependent phosphorylation of STING

Cells were collected in an ice-cold buffer (50 mM Tris-HCl pH 7.4, 100 mM NaCl, 1 mM EGTA, 2 mM DTT, 200 mM sucrose) containing protease inhibitors (25955-11, nacalai tesque), and phosphatase inhibitors (8 mM NaF, 12 mM β-glycerophosphate, 1 mM Na_3_VO_4_, 1.2 mM Na_2_MoO_4_, 5 μM cantharidin, and 2 mM imidazole), homogenized with 2 passages through a 27-gauge needle after 6 passages through a 23-gauge needle, and centrifuged at 3,000 x *g* for 5 min at 4°C. The post-nuclear supernatant was overlaid on 10 μL of 2 M sucrose and centrifuged at 100,000 x *g* for 1 h at 4°C. The membrane fractions were resuspended in a buffer (50 mM Tris-HCl pH 7.4, 100 mM NaCl, 1 mM EGTA, 2 mM DTT, 200 mM sucrose, 20 mM MgCl2, protease inhibitors, and phosphatase inhibitors), and incubated with ATP (1 mM) and recombinant TBK1 (100 ng) at 37°C for 30 min. The samples were then subjected to SDS-PAGE and phosphorylation of STING was detected by western blot.

Western blot. Proteins were separated in polyacrylamide gel and then transferred to polyvinylidene difluoride membranes (Millipore). These membranes were incubated with primary antibodies, followed by secondary antibodies conjugated to horseradish peroxidase. The proteins were visualized by enhanced chemiluminescence using a Fusion SOLO.7S.EDGE (Vilber-Lourmat).

## Acknowledgements

We thank M. Fukuda (Tohoku University) and T. Matsui (Tohoku University) for assistance with ultracentrifugation. This work was supported by JSPS KAKENHI Grant Numbers JP19H00974 (T.T.), JP15H05903 (T.T.), JP17H06164 (H.A.), JP17H06418 (H.A.), JP20H05307 (K.M.), JP20H03202 (K.M.); AMED-PRIME (17939604) (T.T.); JST Center of Innovation program from Japan (JPMJCE1303) (K.M.); The Mitsubishi Foundation (K.M.), the Subsidy for Interdisciplinary Study and Research concerning COVID-19 (Mitsubishi Foundation) (T.T.), Takeda Science Foundation (K.M.), MSD Life Science Foundation (Public Interest Incorporated Foundation) (K.M.), Daiichi Sankyo Foundation of Life Science (K.M.), the Research Foundation For Pharmaceutical Sciences (K.M.), Young Investigator Grant (Graduate School of Life Sciences, Tohoku University) (K.M.), Tokyo Biochemical Research Foundation (K.M.), Grant for Basic Science Research Projects from the Sumitomo Foundation (K.M.), Koyanagi-Foundation (K.M.), and the Nakatomi Foundation (K.M.).

## Author contribution

K.T. designed and performed the experiments, analyzed the data, interpreted the results, and wrote the paper; T.N. designed the experiments with sphingomyelinase; E.O. and F.K. generated TBK1-KO MEFs; Y.N. and M.S. prepared Tag-free recombinant TBK1. H.A. discussed the results; K.M. and T.T. designed the experiments, interpreted the results, and wrote the paper.

## Competing interests

M.S. is a Chief Scientific Officer of Carna Biosciences, Inc. and owns stocks of Carna Biosciences, Inc. T.T. receives research funding from Carna Biosciences, Inc.

## References

1. Palm, N. W. & Medzhitov, R. Pattern recognition receptors and control of adaptive immunity. Immunol Rev 227, 221–233 (2009).

2. Takeuchi, O. & Akira, S. Pattern recognition receptors and inflammation. Cell 140, 805–820 (2010).

3. Wu, J. & Chen, Z. J. Innate immune sensing and signaling of cytosolic nucleic acids. Annu Rev Immunol 32, 461–488 (2014).

4. Sun, L., Wu, J., Du, F., Chen, X. & Chen, Z. J. Cyclic GMP-AMP synthase is a cytosolic DNA sensor that activates the type I interferon pathway. Science 339, 786–791 (2013).

5. Wu, J. et al. Cyclic GMP-AMP is an endogenous second messenger in innate immune signaling by cytosolic DNA. Science 339, 826–830 (2013).

6. Ishikawa, H. & Barber, G. N. STING is an endoplasmic reticulum adaptor that facilitates innate immune signalling. Nature 455, 674–678 (2008).

7. Zhong, B. et al. The adaptor protein MITA links virus-sensing receptors to IRF3 transcription factor activation. Immunity 29, 538–550 (2008).

8. Sun, W. et al. ERIS, an endoplasmic reticulum IFN stimulator, activates innate immune signaling through dimerization. Proc Natl Acad Sci U S A 106, 8653–8658 (2009).

9. Jin, L. et al. MPYS, a novel membrane tetraspanner, is associated with major histocompatibility complex class II and mediates transduction of apoptotic signals. Mol Cell Biol 28, 5014–5026 (2008).

10. Zhang, C. et al. Structural basis of STING binding with and phosphorylation by TBK1. Nature (2019).

11. Zhao, B. et al. A conserved PLPLRT/SD motif of STING mediates the recruitment and activation of TBK1. Nature 569, 718–722 (2019).

12. Tanaka, Y. & Chen, Z. J. STING Specifies IRF3 Phosphorylation by TBK1 in the Cytosolic DNA Signaling Pathway. Sci Signal 5, ra20 (2012).

13. Liu, S. et al. Phosphorylation of innate immune adaptor proteins MAVS, STING, and TRIF induces IRF3 activation. Science 347, aaa2630 (2015).

14. Ogawa, E., Mukai, K., Saito, K., Arai, H. & Taguchi, T. The binding of TBK1 to STING requires exocytic membrane traffic from the ER. Biochem Biophys Res Commun 503, 138–145 (2018).

15. Gui, X. et al. Autophagy induction via STING trafficking is a primordial function of the cGAS pathway. Nature 567, 262–266 (2019).

16. Ran, Y. et al. YIPF5 Is Essential for Innate Immunity to DNA Virus and Facilitates COPII-Dependent STING Trafficking. J Immunol 203, 1560–1570 (2019).

17. Sun, M. S. et al. TMED2 Potentiates Cellular IFN Responses to DNA Viruses by Reinforcing MITA Dimerization and Facilitating Its Trafficking. Cell Rep 25, 3086–3098.e3 (2018).

18. Liu, Y. et al. Activated STING in a vascular and pulmonary syndrome. N Engl J Med 371, 507–518 (2014).

19. Jeremiah, N. et al. Inherited STING-activating mutation underlies a familial inflammatory syndrome with lupus-like manifestations. J Clin Invest 124, 5516–5520 (2014).

20. Watkin, L. B. et al. COPA mutations impair ER-Golgi transport and cause hereditary autoimmune-mediated lung disease and arthritis. Nat Genet 47, 654–660 (2015).

21. Mukai, K. et al. Activation of STING requires palmitoylation at the Golgi. Nat Commun 7, 11932 (2016).

22. Dobbs, N. et al. STING Activation by Translocation from the ER Is Associated with Infection and Autoinflammatory Disease. Cell Host Microbe 18, 157–168 (2015).

23. Deng, Z. et al. A defect in COPI-mediated transport of STING causes immune dysregulation in COPA syndrome. J Exp Med 217, e20201045 (2020).

24. Mukai, K. et al. Homeostatic regulation of STING by retrograde membrane traffic to the ER. Nat Commun 12, 61 (2021).

25. Lepelley, A. et al. Mutations in COPA lead to abnormal trafficking of STING to the Golgi and interferon signaling. J Exp Med 217, (2020).

26. Taguchi, T. & Mukai, K. Innate immunity signalling and membrane trafficking. Curr Opin Cell Biol 59, 1–7 (2019).

27. Hansen, A. L. et al. Nitro-fatty acids are formed in response to virus infection and are potent inhibitors of STING palmitoylation and signaling. Proc Natl Acad Sci U S A 115, E7768–E7775 (2018).

28. Hansen, A. L., Mukai, K., Schopfer, F. J., Taguchi, T. & Holm, C. K. STING palmitoylation as a therapeutic target. Cell Mol Immunol 16, 236–241 (2019).

29. Ishikawa, H., Ma, Z. & Barber, G. N. STING regulates intracellular DNA-mediated, type I interferon-dependent innate immunity. Nature 461, 788–792 (2009).

30. Lippincott-Schwartz, J. et al. Microtubule-dependent retrograde transport of proteins into the ER in the presence of brefeldin A suggests an ER recycling pathway. Cell 60, 821–836 (1990).

31. Duran, J. M. et al. Sphingomyelin organization is required for vesicle biogenesis at the Golgi complex. EMBO J 31, 4535–4546 (2012).

32. Linder, M. E. & Deschenes, R. J. Palmitoylation: policing protein stability and traffic. Nat Rev Mol Cell Biol 8, 74–84 (2007).

33. Nishimura, T. et al. Oxysterol-binding protein (OSBP) is required for the perinuclear localization of intra-Golgi v-SNAREs. Mol Biol Cell 24, 3534–3544 (2013).

34. Pol, A. et al. Cholesterol and fatty acids regulate dynamic caveolin trafficking through the Golgi complex and between the cell surface and lipid bodies. Mol Biol Cell 16, 2091–2105 (2005).

