## Supplementary Figures for "Implications of cholesterol and sphingomyelin in STING phosphorylation by TBK1"

### supplementary figure 1

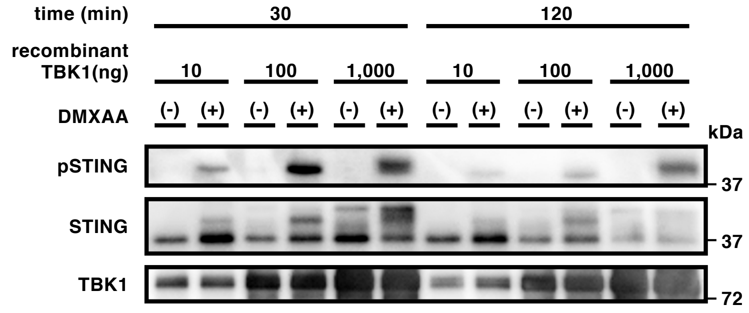


**supplementary figure 1** | supplementary materials related to Fig. 2.
TBK1-KO MEFs were stimulated with DMXAA (25 μg/ml) for 0 or 1 h, homogenized in isotonic buffer, and centrifuged at 3,000 x *g* for 5 min. The resulting post-nuclear supernatants were then centrifuged at 100,000 x *g* for 1 h, and the pellets were resuspended in isotonic buffer. The resuspended membrane fractions were incubated with ATP and recombinant TBK1 (10 ng, 100 ng, or 1,000 ng) at 37 °C for 30 or 120 min.

### supplementary figure 2

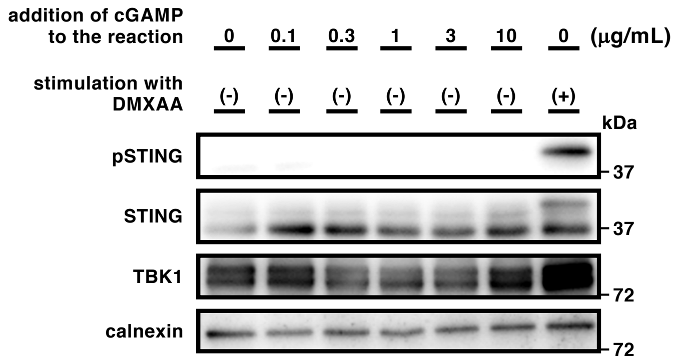


**supplementary figure 2** | supplementary materials related to Fig. 3b.
Post-nuclear supernatants of unstimulated TBK1-KO MEFs were centrifuged at 100,000 x *g*. The resulting membrane fraction was resuspended and incubated with recombinant TBK1 and ATP in the presence of 2’3’-cGAMP (0.1 μg/mL - 10 μg/mL) at 37 °C for 30 min. Phosphorylation of STING at Ser365 was examined by western blot. Microsomal membrane fraction prepared from DMXAA-stimulated cells was used as a positive control for the reaction.
